# Elucidation of cryptic and allosteric pockets within the SARS-CoV-2 protease

**DOI:** 10.1101/2020.07.23.218784

**Authors:** Terra Sztain, Rommie Amaro, J. Andrew McCammon

## Abstract

The SARS-CoV-2 pandemic has rapidly spread across the globe, posing an urgent health concern. Many quests to computationally identify treatments against the virus rely on *in silico* small molecule docking to experimentally determined structures of viral proteins. One limit to these approaches is that protein dynamics are often unaccounted for, leading to overlooking transient, druggable conformational states. Using Gaussian accelerated molecular dynamics to enhance sampling of conformational space, we identified cryptic pockets within the SARS-CoV-2 main protease, including some within regions far from the active site and assed their druggability. These pockets can aid in virtual screening efforts to identify a protease inhibitor for the treatment of COVID-19.

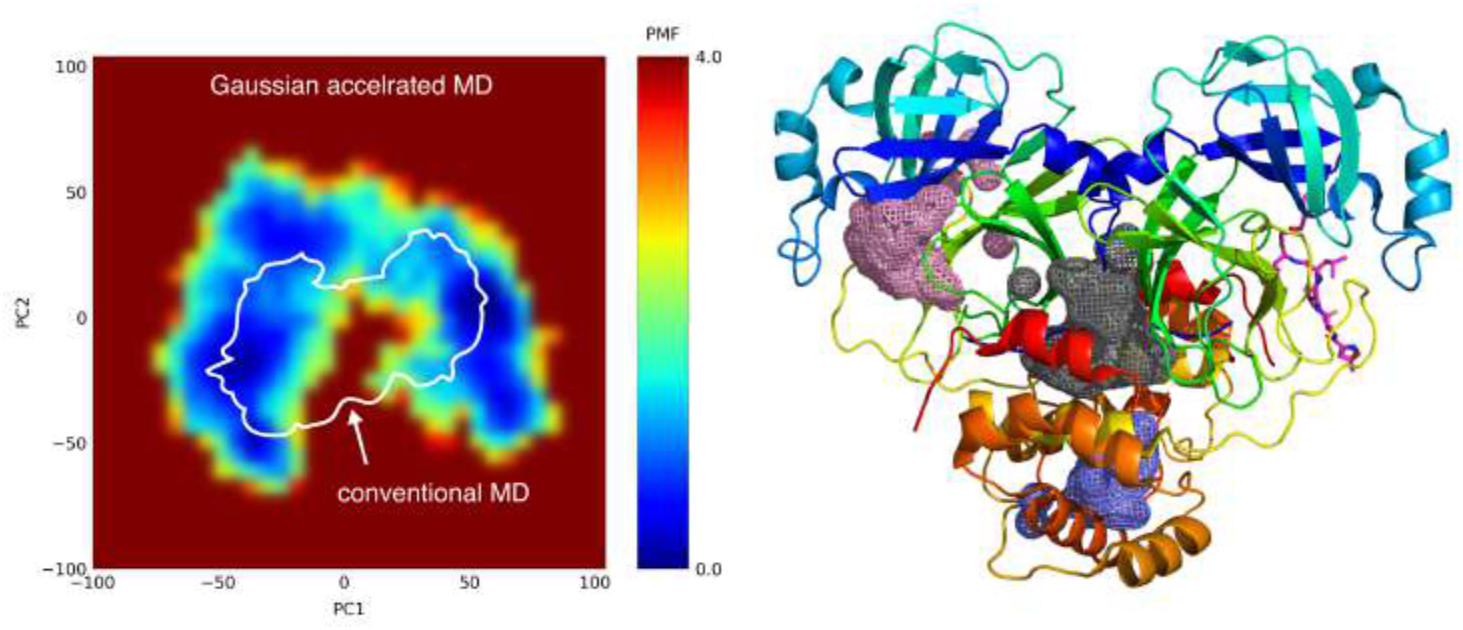

## Introduction

In December 2019, the World Health Organization learned of a novel coronavirus which has since rapidly spread, leading to a global pandemic. The SARS-CoV-2 virus, causing the disease named COVID-19 has reached 15 million cases, taking over 600,000 lives as of July 22, 2020.^1^ The genetic and structural similarity of the SARS-CoV-2 virus to the agents of the severe acute respiratory syndrome coronavirus (SARS-CoV) epidemic in 2003 and Middle East respiratory syndrome coronavirus (MERS-CoV) epidemic in 2012 provide a basis of information for understanding and ultimately treating or preventing this disease, however no highly effective treatment or vaccine exists for any human infecting coronavirus.

Proteases are responsible for activating viral proteins for particle assembly and have proved successful targets for antiviral agents; most notable are the protease inhibitors used to treat HIV and Hepatitis C.^2,3^ The main protease of SARS-CoV-2, called M^pro^ or 3CL^pro^ was the first SARS-CoV-2 protein deposited to the protein databank (PDB) on January 26^th^ 2020.^4^ This structure was crystalized with a covalent inhibitor (N3) identified from computer aided drug design, and validated biochemically. Currently, over one hundred M^pro^ structures exist in the PDB and massive efforts to discover a successful inhibitor are underway.^4,5,6,7^ Many such efforts involve high throughput virtual screening, which exploits the power of computational docking to screen millions of molecules *in silico* to narrow down a few “hits” for lead optimization.

Incorporation of molecular dynamics (MD) has significantly improved the ability to identify promising protein inhibitors.^8^ Docking to protein ensembles obtained from MD simulations is often employed to consider multiple target states that remain elusive in static crystal structures.^9^ Conventional MD simulations are however limited in the amount of conformational space that can be sampled due to the amount of time required to traverse energy barriers between stable conformational states.^10^ In the present study, we used the enhanced sampling technique, Gaussian accelerated MD (GaMD),^11^ to overcome such barriers. GaMD adds a harmonic boost to the potential energy up to a threshold, effectively “filling in the wells” creating a smoother potential energy surface. This allowed extensive conformational sampling of the SARS-CoV-2 main protease. High throughput virtual screening (HTVS) of almost 72,000 compounds to an ensemble of over 80 of these structures revealed various molecular architectures predicted to bind to the SARS-CoV-2 protease, which can serve as a basis for structure activity relationship studies to identify an inhibitor for the treatment of COVID-19. These studies also revealed that a distal site and the dimer interface, in addition to the active site, may serve as viable targets for inhibitor development.

## Results

GaMD simulations were performed on four systems built from the first published M^pro^ structure PDB 6LU7.^4^ Both a monomer and dimer were simulated in the presence and absence of the co-crystalized N3 inhibitor. Five independent simulations of 200 ns, for an aggregate of 1μs were performed for each system. Three regions were defined to investigate potentially druggable pockets. The active site lies in the N-terminal top lobe of the heart shaped protease. The C-terminal region was examined as a potential allosteric site, in addition to the dimer interface region (Figure 1A). POVME^12^ was used to calculate the pocket volumes of one hundred distinct frames from each system, obtained from clustering based on root mean squared deviation (RMSD) from the first frame. An array of pocket volumes was sampled between 20 and 180 Å^3^ for the active site pocket, starting from 170 Å^3^ in the crystal structure. The interface spanned from 100 to 420 Å^3^ starting from 209 Å^3^, and the distal site from 0 to 80 Å^3^ starting from 18 Å^3^ (Figure 1B-D). Simulations with the N3 ligand present sampled larger mean volumes than the apo simulations, however the average dimer interface region was greater in the apo simulation (Figure S1). Trajectory visualization (Movie S1) and overlay of several pockets revealed loop dynamics leading to distinct conformations (Figure 2) which were used for an ensemble docking HTVS approach, and further analyzed for druggability using the PockDrug webserver.^13^

**Figure 1.**
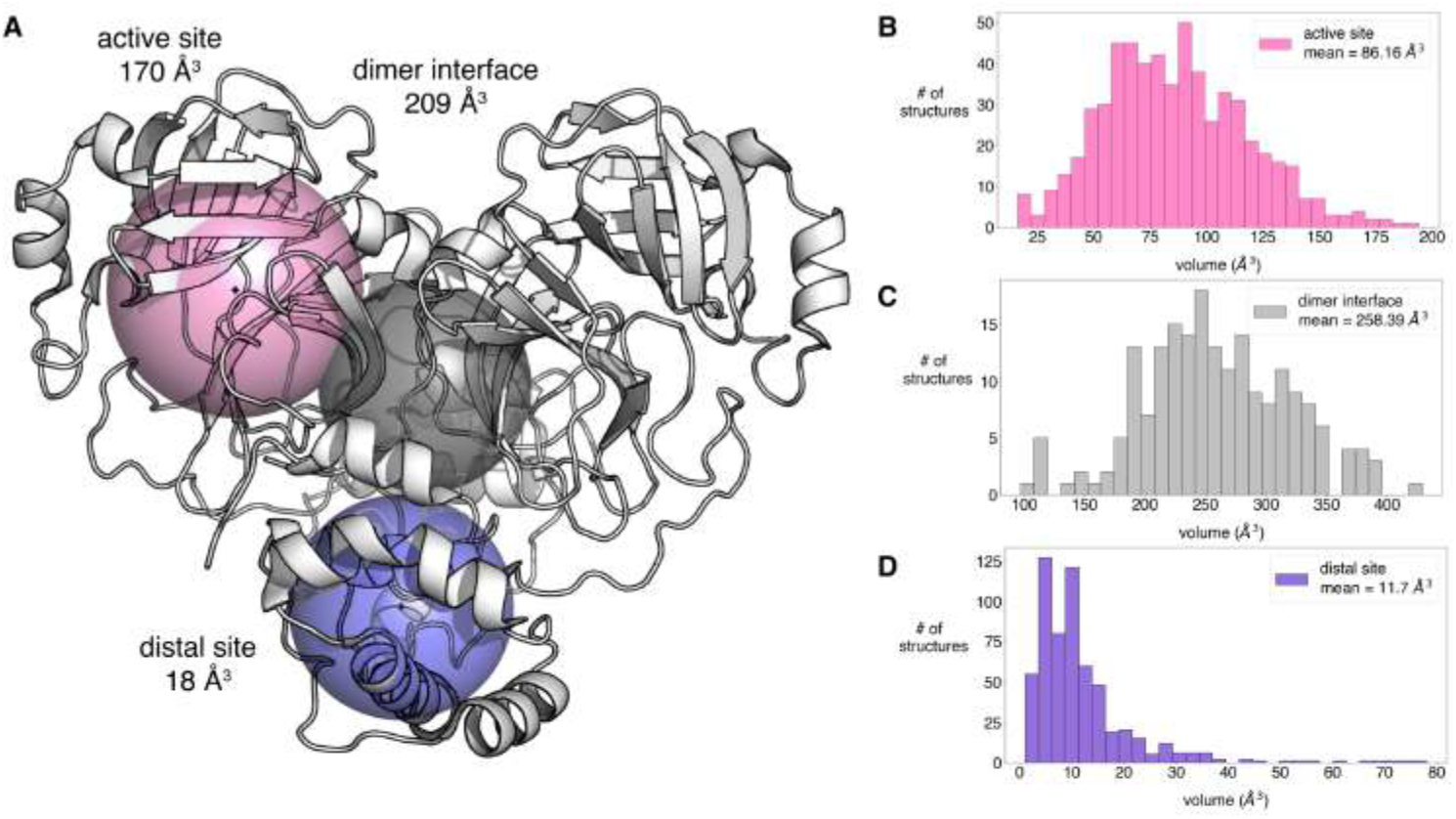
Pocket analysis of GaMD simulations of M^pro^. (A) Active site, dimer interface, and distal site definitions for pocket calculations. The inclusion sphere was centered on the center of mass (COM) of the top and bottom protein domains, for active and distal site with 12 Å and 10 Å radii respectively. The dimer interface center was defined as the COM of the residues within 3.5 Å of the other protein chain for both chains. A 10 Å sphere was used for the dimer interface calculation. (B-D) Histograms showing the distribution of pocket volumes calculated from the aggregate 4 μs GaMD simulations. Each of the four systems simulated were narrowed down to 100 frames by clustering based on all atom RMSD from the first frame. The dimer simulations provided 2 data points for the active and distal site regions and one point for the interface region, while the monomer simulations did not contain a dimer interface.

**Figure 2.**
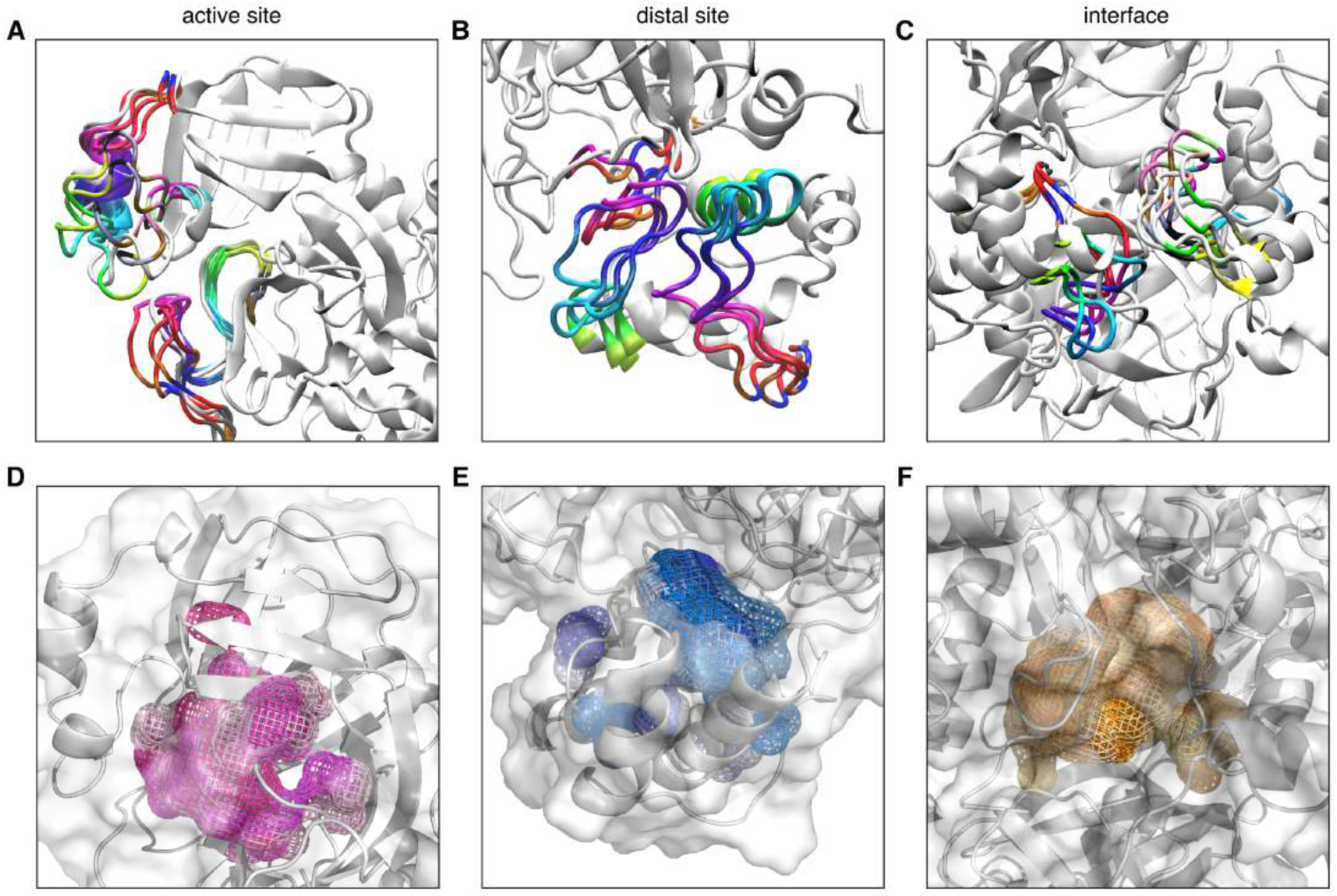
Cryptic pockets identified from GaMD simulations. (A-C) Overlay of four frames from dimer simulation highlighting loop dynamics leading to various pocket conformations. (D-F) Overlay of six pockets shown outlined in mesh, colored in varying shades of pink for the active site, blue for the distal site, and orange for the interface.

The identified pockets were clustered into five structures for each region and the largest pocket volume was also included for a total of 82 structures subject to HTVS of 71,789 compounds from the ChemBridge CombiSet diversity library. A total of 14,340 unique compounds passed the Glide^14^ docking funnel and XP score filter, including 6303 docking to the active site, 8598 docking to the distal site, and 9538 docking to the dimer interface compared to 13, 48 and 1710 respectively, docking to the crystal structure. The top compounds which docked to the dimer interface gave calculated Glide scores around −10, compared to −9 for the active site region, and −8 for the distal region, while the compounds docked to the crystal structure scored around −8, −6, and −3, respectively with larger negative scores indicating lower calculated energy, or greater docking affinity (Table S1-4, molecule identifiers removed for pre-print). Certain pockets showed significantly higher affinity for small molecule docking upon HTVS. For example, the top 10 overall scoring compounds which docked to the active site were all docked to cluster number 39 from the dimer simulation which was simulated with N3 bound in the pocket and has a calculated pocket volume of 160 Å^3^. N3 was also docked to each of the structures as a control, however it scored poorly with the general docking protocol which is not optimized for considering covalent bonding. The significant increase in identified molecules, and better docking score values obtained from GaMD simulations compared to the crystal structure highlights the value of sampling increased conformational space for identifying potential protein inhibitors.

The druggability probability for each structure subjected to HTVS was calculated based on a variety of factors including geometry, hydrophobicity, aromaticity and others by the PockDrug webserver^13^ (Table S1-4). The crystal structure active site region has predicted druggability of .68 ± .07, while the dimer interface and distal site scored better, .91 ± .05 and .90 ± .05 respectively. The predicted druggability was not correlated to the pockets with the best small molecule docking scores, indicating several factors should be considered in evaluating the identified conformations. For example cluster 39 discussed above had a calculated druggability probability of .42 ± .05. Cluster 0 from the apo dimer simulation gave a predicted druggability of 1.0 ± 00 for the active site region. Four clusters also gave a perfect probability for the distal site region, and the highest probability for the interface region was .93 ± .02 (Table S1-4).

The comparable number of total molecules, value of docking scores, and predicted druggability of the distal site and dimer interface indicate that these regions may be viable targets for inhibitor development in addition to the active site. Dimer association is necessary for catalytic activity of the protease,^4^ therefore it is reasonable to assume binding of a molecule which prevents this association would inactivate the protease. The significance of the distal regions is less well understood, though computational investigations support a potential allosteric role.^17,18^ To study this further, we examined the correlated motions of each residue throughout the simulations.

In the monomer simulation, the active site region was positively correlated to the distal site region, and negatively correlated to the region that connects the two, whereas the inverse was observed in the dimer simulation (Figure 3, S2,3). These correlations resulted in primary motions detected by principal component analysis comprising of more hinge-like in the monomer versus twisting in the dimer (Figure S4). These motions were similar in both the apo and N3 bound simulations. In the dimer simulations, the active site of chain A was slightly correlated to the active site of chain B, and the distal regions were highly anticorrelated, twisting in opposite directions. (Figure S4). Regardless of positive or negative correlation values, the distal and active site show a dynamic interdependence on each other in both the apo and dimer forms.

**Figure 3.**
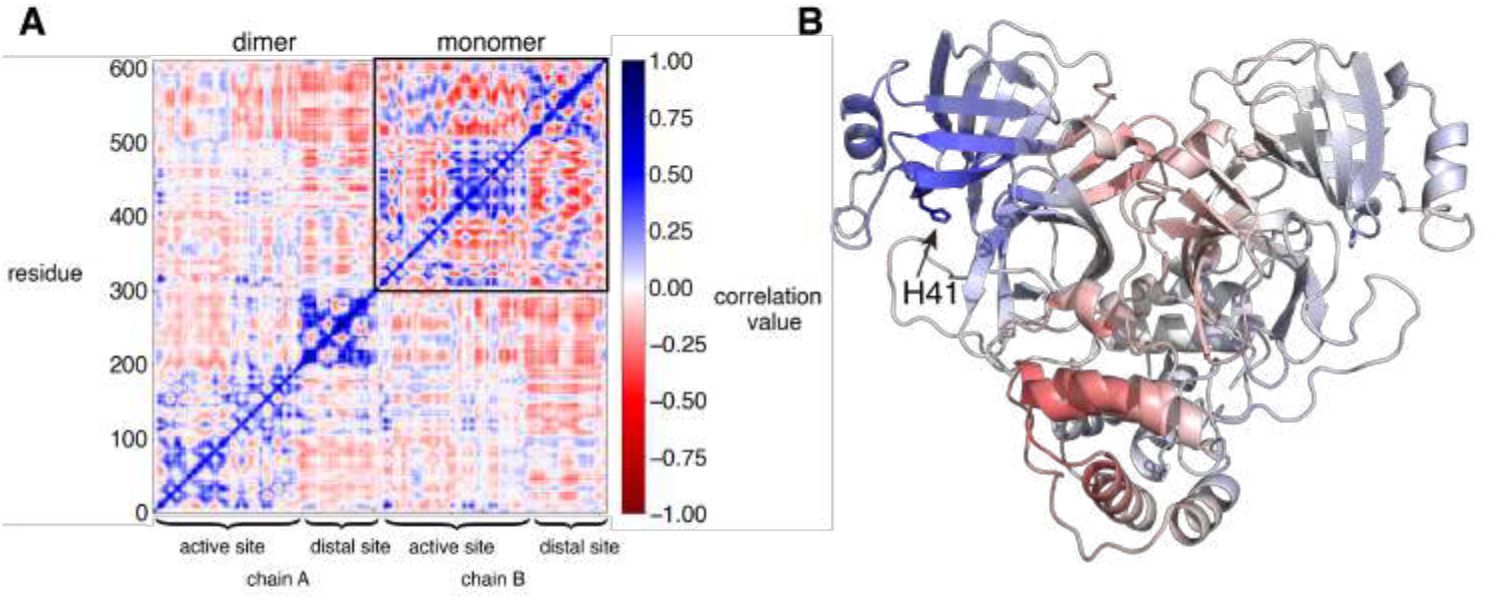
Correlated motions of each residue. (A) Correlation matrix calculated from aggregate 1 μs simulation of apo M^pro^. Calculation from the monomer simulation is shown as an inset in the upper right corner of the calculation from the dimer simulation. (B) Structural representation of correlated motions, with each residue colored based on correlation to the catalytic histidine.

## Conclusion

The enhanced conformational space obtained by Gaussian accelerated molecular dynamics simulations of the SARS-CoV-2 main protease, M^pro^, revealed cryptic pockets not detectable from the crystal structure more efficiently than brute force simulations. High throughput virtual screening to these pockets allowed identification of two orders of magnitude greater number of hit molecules than the crystal structure to the active site region, and almost an order of magnitude greater total molecules to all regions investigated. Both the dimer interface and the distal site region of M^pro^ formed favorable interactions *in silico* with the drug-like molecules. The docking scores and predicted druggability for each region screened were comparable to that of the active site. Correlated motions of residues at each site indicate that binding to one site could be translated to the active site region in an allosteric manner. The pockets identified here, in addition to the microsecond simulations carried out by DE Shaw Research^19^ and millisecond simulations through Folding@home^20^, can serve as a basis for further docking of potential drug molecules, which can contribute towards development of an M^pro^ inhibitor for the treatment of COVID-19.

## Data Accession

Data from this study available from the corresponding author, upon reasonable request.

## Methods

### Simulation preparation

The first crystal structure of SARS-CoV-2 main protease, PDB ID 6LU7^4^ was used as a starting point for all simulations. This structure was crystalized with the inhibitor N3. For apo simulations, the inhibitor was deleted from the PDB file. For dimer simulations, the PyMOL^15^ symexp command was used to generate initial coordinates for the second chain based on symmetry. Four systems were simulated: apo-monomer, apo-dimer, holo-monomer and holo-dimer (N3 bound to chain B). Protonation states of titratable residues were determined using the H++ webserver^21^ with 0.15 M salinity, 10 internal dielectric constant, 80 external dielectric constant, and pH 7.4. Histidine 64, 80, and 164 were thus named HID to indicate delta nitrogen protonation before protonation of entire protein using tleap from AmberTools 18.^22^ The ff14SB^23^ forcefield was used for proteogenic residues. For simulations with the N3 bound, the antechamber package of AmberTools 18 was used to generate GAFF^24^ forcefield parameters for N3. Gaussian 09 was used to calculate partial charges of atoms according to the RESP^25^ method with HF/6-31G* level of theory. A TIP3P isometric water box was added with at least 10 Å buffer between solute and edges of the box. Enough Na+ was added to neutralize the system, then Na+ and Cl- ions were added to a final concentration of 150 mM. Minimization was carried out in two steps. First the solute was restrained with a 500 kcal / mol restraint force to minimize the solvent for 10,000 cycles, followed by unrestrained minimization for 300,000 cycles. Next the system was heated to 310 K over 350 ps using an isothermal-isovolumetric (NVT) ensemble, followed by isothermal-isobaric (NPT) equilibration for 1 ns.

### Gaussian accelerated molecular dynamics

Gaussian accelerated molecular dynamics (GaMD)^11^ pmemd.cuda implementation of Amber 18 was used to generate 5 independent trajectories of 200 ns, an aggregate of 1 μs for each system. The dual boost method was employed, adding a biasing force to both the total and dihedral potential energy. The threshold energy was set to the upper bound. GaMD production of equilibrated systems was carried out in 5 stages. First, 200 ps of preparatory conventional MD simulation were carried out, without any statistics collected. Second, 1 ns of conventional MD was carried out to collect potential statistics V_max_, V_min_, V_avg_, and σV. Next, 800 ps of GaMD were carried out with a boost potential applied with fixed parameters. Then, 50 ns of GaMD were carried out with updated boost parameters and finally 150 ns of GaMD were carried out with fixed boost parameters. All production simulations were carried out using NVT, with periodic boundary conditions and 2 fs timesteps. The SHAKE^26^ algorithm was used to restrain non polar hydrogen bonds and TIP3P water molecules. Particle Mesh Ewald^27^ method was used for electrostatic interactions with a 10 Å cutoff for nonbonded interactions. Langevin thermostat was used for temperature regulation with a collision frequency of 5 ps^−1^.

### Pocket analysis

For each system simulated, the 1 μs of aggregated GaMD simulation time was clustered based on the RMSD from the first frame using cpptraj^28^ hierarchical algorithm to obtain 100 distinct structures for pocket analysis. Three regions were defined for individual calculations: the active site pocket, distal site pocket, and dimer interface. Pocket volume was calculated using POVME 2^12^ with 1 Å grid spacing. The point inclusion sphere was determined based on the average center of mass of residues 7-198 with a 12 Å radius for the active site pocket, average center of mass of residues 198-306 with a 10 Å radius for the distal site pocket, and the average center of mass between residues within 3.5 Å of other dimer with a 10 Å radius for the dimer interface. Pocket volumes were used to cluster pocket regions for high throughput virtual screening. The Agglomerative Clustering module of Scikit Learn^29^ was used to generate 5 clusters of data points, and the structure with pocket volume closest to the cluster center was subject to high throughput virtual screening (HTVS). The largest pocket volume of each region for each system, was also subject to HTVS.

### High throughput virtual screening

Schrödinger Virtual Screening Workflow was followed using Schrödinger release 2020-1.^14^ ChemBridge CombiSet Library of 71,789 compounds were prepared using LigPrep with empirical pKa prediction (Epik).^30^ Protein structures were prepared with the prepwizard module, after removing the N3 ligand if present. Glide was used to generate a docking grid, centered on the same center that was used in pocket volume calculations, with a 10 Å^3^ inner box and a 30 Å^3^ outer box. The inner box defines the volume which the ligand center can explore, while the outer box defines the volume within which grid potentials are calculated. The vsw module was used to perform HTVS, followed by SP docking and XP docking, eliminating 70% of the lowest scoring compounds at each step. Glide XP docking was performed on the remaining models for final scoring and analysis.

### Additional analysis

For the graphical abstract, to demonstrate enhanced sampling of conformational space, reweighting of the potential energy was carried out on one 200 ns simulation of the N3 bound dimer using PyReweighting toolkit.^31^ Reweighting was carried out using the Maclaurin series expansion to the 10^th^ order with a bin size of 6 and maximum energy of 100, based on the first two principal components obtained from projection of displacement vectors of each of the backbone atoms onto a diagonalized mass-weighted covariance matrix after rms fitting of every atom except protons to the first frame, calculated using cpptraj as described previously.^32^ One conventional 200 ns MD simulation was performed for comparison and outline in the figure. Correlation matrices for each residue were obtained from cpptraj after rms fitting to all atoms of the first frame. Principal motions were defined based on visualization of the first normal mode using the Normal Mode Wizard plugin of VMD^33^ calculated from projection of displacement vectors of each of the backbone atoms onto a diagonalized mass-weighted covariance matrix after rms fitting of every atom except protons to the first frame for 1 μs aggregate simulation of each system.

## Acknowledgments

We thank Michael D. Burkart for advisement, and the NSF GRFP for supporting T.S. under grant number DGE-1650112. Simulations were carried out using GPU nodes of the Triton Shared Computing Cluster (TSCC) at the San Diego Supercomputer Center (SDSC). This work was supported by NIH GM031749.

