## Supplementary Information for "Elucidation of cryptic and allosteric pockets within the SARS-CoV-2 protease"

**Supplementary Figures**

**Figure S1** Pocket volume of each region in apo versus holo simulations

**Figure S2** Correlation matrix calculated from aggregate 1 μs of each simulation

**Figure S3** Structural representation of correlated motions to the catalytic histidine.

**Figure S4** Principal motions during dimer and monomer simulations

**Supplementary Tables**

**Table S1** Top scores and druggability for each of the GaMD structures used in the HTVS

**Table S2** Top 10 scores and druggability overall for the active site region

**Table S3** Top 10 scores and druggability overall for the distal site region

**Table S4** Top 10 scores and druggability overall for the dimer interface region

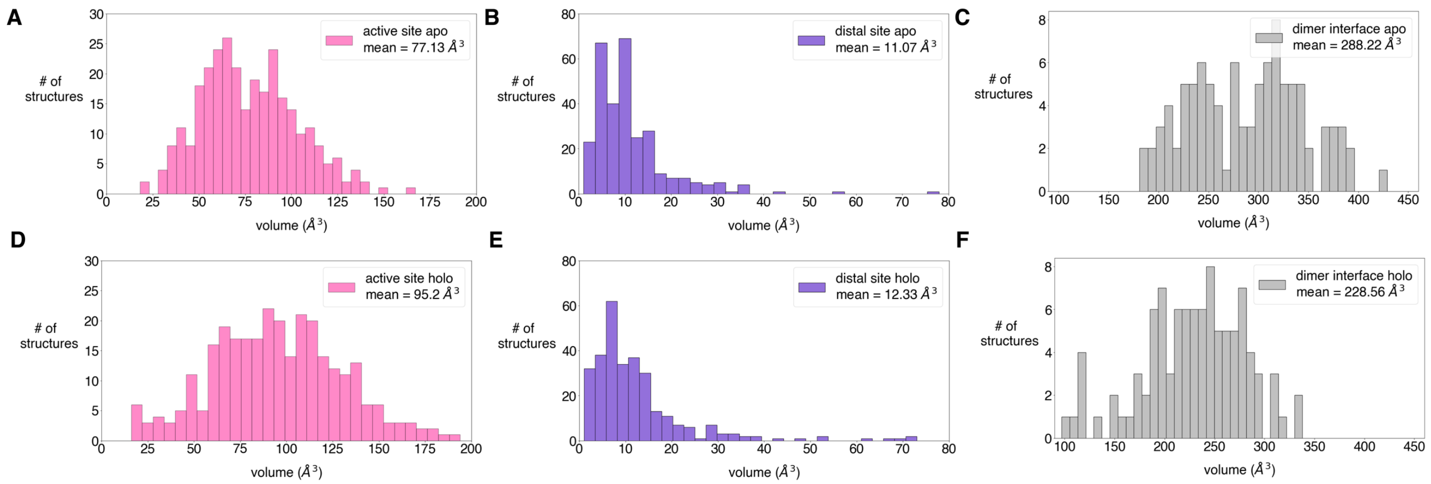


**Figure S1** Pocket volume of each region in apo versus holo simulations. Histograms show average pocket volume calculated from aggregate 1 µs dimer and monomer simulations of $M^{pro}$ in the apo (A-C) and holo, N3 bound state (D-F).

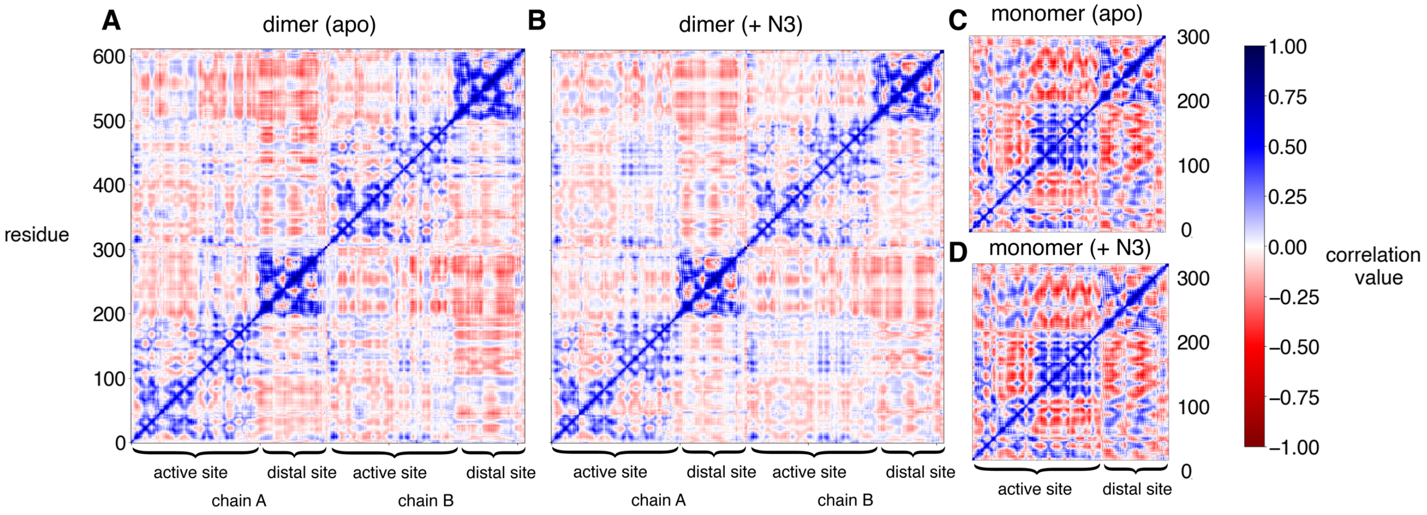


**Figure S2** Correlation matrix calculated from aggregate 1 μs of each simulation. (A) Apo $M^{pro}$ dimer (B) $M^{pro}$ dimer with N3 bound to chain B. (C) Apo $M^{pro}$ monomer. (D) $M^{pro}$ monomer with N3 bound.

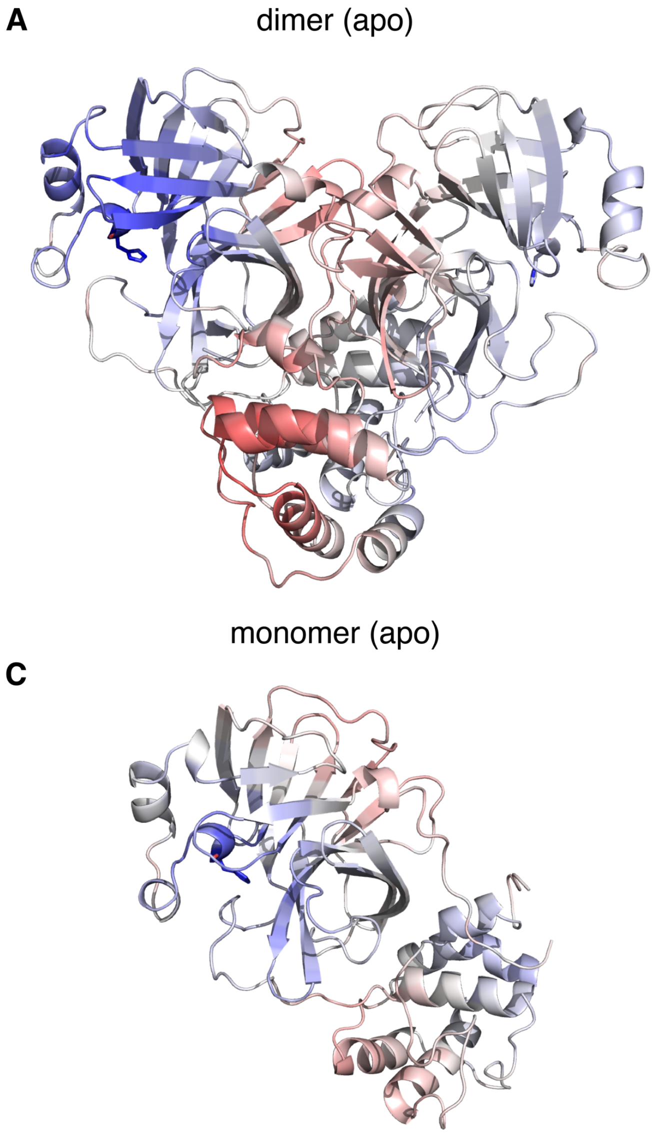


**Figure S3** Structural representation of correlated motions to the catalytic histidine. Each residue is colored based on correlation value to the catalytic histidine residue 41 for (A) Apo $M^{pro}$ dimer, (B) $M^{pro}$ dimer with N3 bound to chain B, (C) Apo $M^{pro}$ monomer, and (D) $M^{pro}$ monomer with N3 bound.

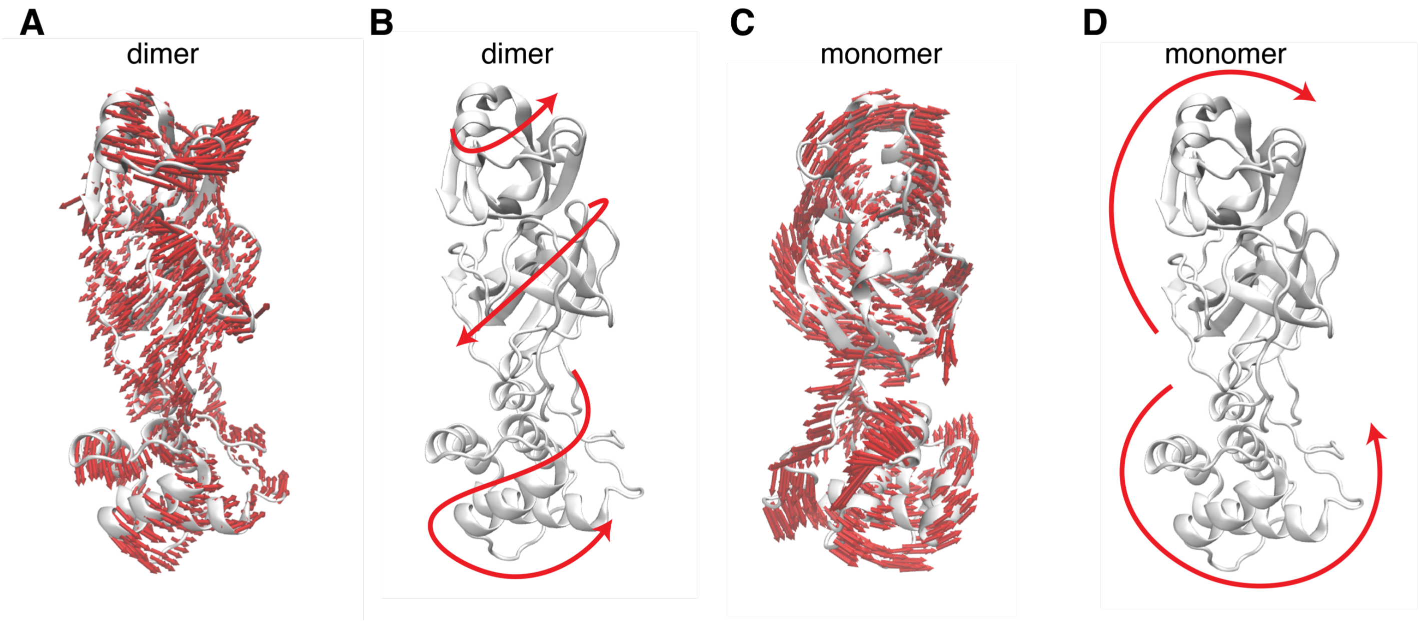


**Figure S4** Principal motions during dimer and monomer simulations. The normal modes from the first principal component were projected onto one monomer from the dimer simulation (A) and monomer simulation (C) using VMD. The resulting overall "twisting" motion observed in the dimer is schematically shown by the red arrows in (B) and the "hinging" motion observed in the monomer is shown in (D).

**Table S1** Top scoring compound for each of the GaMD structures used in the HTVS

| simulation | region (chain) | rms cluster | Gscore[a] | drugability[b] |
|---|---|---|---|---|
| monomer | active_site | 94 | -5.11 | .86 ± .02 |
| monomer | active_site | 65 | -2.98 | .75 ± .06 |
| monomer | active_site | 0 | -5.11 | .80 ± .07 |
| monomer | active_site | 50 | -4.14 | .97 ± 00 |
| monomer | active_site | 98 | -4.94 | .90 ± .01 |
| monomer | distal_site | 19 | -5.39 | 1.0 ± 00 |
| monomer | distal_site | 3 | -6.23 | .81 ± .07 |
| monomer | distal_site | 62 | -5.03 | 1.0 ± 00 |
| monomer | distal_site | 0 | -6.55 | .98 ± .01 |
| monomer | distal_site | 5 | -5.47 | .99 ± 00 |
| monomer | distal_site | 61 | -6.84 | 1.0 ± 00 |
| monomer_apo | active_site | 97 | -4.07 | .27 ± .06 |
| monomer_apo | active_site | 1 | -3.83 | .99 ± 00 |
| monomer_apo | active_site | 23 | -2.87 | .98 ± 00 |
| monomer_apo | active_site | 17 | -3.92 | .94 ± 00 |
| monomer_apo | active_site | 72 | -8.33 | .72 ± .02 |
| monomer_apo | distal_site | 5 | -4.06 | .82 ± .06 |
| monomer_apo | distal_site | 7 | -4.99 | .80 ± .01 |
| monomer_apo | distal_site | 9 | -4.98 | .62 ± .02 |
| monomer_apo | distal_site | 15 | -4.73 | .26 ± .13 |
| monomer_apo | distal_site | 23 | -3.67 | .98 ± .02 |
| monomer_apo | distal_site | 58 | -7.06 | .99 ± 00 |
| dimer | active_site_A | 56 | -4.59 | .69 ± .04 |
| dimer | active_site_A | 31 | -4.74 | .90 ± .05 |
| dimer | active_site_A | 43 | -7.64 | .97 ± .01 |
| dimer | active_site_A | 20 | -6.72 | .48 ± .08 |
| dimer | active_site_A | 42 | -8.26 | .86 ± .03 |

| | | | | |
|---|---|---|---|---|
| dimer | active_site_A | 30 | -8.09 | .81 ± .01 |
| dimer | active_site_B | 15 | -6.87 | .58 ± .13 |
| dimer | active_site_B | 0 | -8.37 | .66 ± .05 |
| dimer | active_site_B | 10 | -7.73 | .86 ± .03 |
| dimer | active_site_B | 93 | -4.94 | .78 ± .02 |
| dimer | active_site_B | 99 | -5.89 | .78 ± .09 |
| dimer | active_site_B | 39 | -9.44 | .42 ± .05 |
| dimer | distal_site_A | 9 | -5.21 | .93 ± .03 |
| dimer | distal_site_A | 19 | -5.08 | .56 ± .01 |
| dimer | distal_site_A | 36 | -5.31 | .97 ± .01 |
| dimer | distal_site_A | 67 | -5.79 | .99 ± 00 |
| dimer | distal_site_A | 50 | -5.61 | .45 ± .06 |
| dimer | distal_site_A | 74 | -6.97 | .56 ± .04 |
| dimer | distal_site_B | 13 | -5.3 | 1.0 ± 00 |
| dimer | distal_site_B | 1 | -7.45 | .99 ± 00 |
| dimer | distal_site_B | 89 | -6.84 | .32 ± .03 |
| dimer | distal_site_B | 16 | -7.96 | .82 ± .05 |
| dimer | distal_site_B | 17 | -6.34 | .63 ± .08 |
| dimer | distal_site_B | 4 | -8.09 | .98 ± 00 |
| dimer | interface | 25 | -9.68 | .75 ± .03 |
| dimer | interface | 73 | -7.76 | .93 ± .02 |
| dimer | interface | 6 | -9.3 | .80 ± .02 |
| dimer | interface | 15 | -9.72 | .84 ± .05 |
| dimer | interface | 35 | -8.62 | .74 ± .06 |
| dimer_apo | active_site_A | 14 | -5.22 | .51 ± .05 |
| dimer_apo | active_site_A | 95 | -6.29 | .77 ± .03 |
| dimer_apo | active_site_A | 71 | -6.81 | .75 ± .02 |
| dimer_apo | active_site_B | 28 | -7.2 | .90 ± .01 |
| dimer_apo | active_site_B | 85 | -6.18 | .93 ± .02 |

| | | | | |
|---|---|---|---|---|
| dimer_apo | active_site_B | 62 | -8.26 | .87 ± .04 |
| dimer_apo | active_site_B | 0 | -6.82 | 1.0 ± 00 |
| dimer_apo | active_site_B | 18 | -7.73 | .90 ± .06 |
| dimer_apo | active_site_B | 3 | -6.33 | .79 ± .08 |
| dimer_apo | distal_site_A | 1 | -4.17 | .85 ± .01 |
| dimer_apo | distal_site_A | 76 | -4.4 | .79 ± .02 |
| dimer_apo | distal_site_A | 6 | -5.94 | .98 ± .01 |
| dimer_apo | distal_site_A | 15 | -4.07 | .97 ± .01 |
| dimer_apo | distal_site_A | 35 | -4.68 | .99 ± 00 |
| dimer_apo | distal_site_B | 4 | -7.75 | .93 ± .01 |
| dimer_apo | distal_site_B | 2 | -7.95 | .34 ± .09 |
| dimer_apo | distal_site_B | 31 | -7 | .99 ± 00 |
| dimer_apo | distal_site_B | 26 | -8.65 | .97 ± 00 |
| dimer_apo | distal_site_B | 15 | -7.25 | .99 ± 00 |
| dimer_apo | distal_site_B | 35 | -6.1 | .94 ± .01 |
| dimer_apo | interface | 50 | -8.69 | .53 ± .08 |
| dimer_apo | interface | 89 | -8.39 | .61 ± .06 |
| dimer_apo | interface | 10 | -10.07 | .65 ± .02 |
| dimer_apo | interface | 32 | -8.7 | .92 ± .03 |
| dimer_apo | interface | 39 | -9.99 | .83 ±.03 |
| dimer_apo | interface | 0 | -8.74 | .65 ± .02 |

[a] Schrodinger Glide Gscore

[b] Calculated from PockDrug webserver

**Table S2** Top 10 scoring compounds overall for the active site region

| simulation | region (chain) | rms cluster | Gscore[a] | druggability[b] |
|---|---|---|---|---|
| dimer | active_site_B | 39 | -9.44 | .42 ± .05 |
| dimer | active_site_B | 39 | -9.38 | .42 ± .05 |
| dimer | active_site_B | 39 | -9.18 | .42 ± .05 |
| dimer | active_site_B | 39 | -9.14 | .42 ± .05 |
| dimer | active_site_B | 39 | -9.04 | .42 ± .05 |
| dimer | active_site_B | 39 | -9.03 | .42 ± .05 |
| dimer | active_site_B | 39 | -9.02 | .42 ± .05 |
| dimer | active_site_B | 39 | -8.78 | .42 ± .05 |
| dimer | active_site_B | 39 | -8.77 | .42 ± .05 |
| dimer | active_site_B | 39 | -8.76 | .42 ± .05 |

[a] Schrodinger Glide Gscore

[b] Calculated from PockDrug webserver

**Table S3** Top 10 scoring compounds overall for the distal site region

| simulation | region (chain) | rms cluster | Gscore[a] | druggability[b] |
|---|---|---|---|---|
| dimer_apo | distal_site_B | 26 | -8.65 | .97 ± 00 |
| dimer | distal_site_B | 4 | -8.09 | .98 ± 00 |
| dimer_apo | distal_site_B | 26 | -8.05 | .97 ± 00 |
| dimer | distal_site_B | 16 | -7.96 | .82 ± .05 |
| dimer_apo | distal_site_B | 2 | -7.95 | .34 ± .09 |
| dimer_apo | distal_site_B | 26 | -7.89 | .97 ± 00 |
| dimer_apo | distal_site_B | 26 | -7.86 | .97 ± 00 |
| dimer | distal_site_B | 4 | -7.75 | .98 ± 00 |
| dimer_apo | distal_site_B | 4 | -7.75 | .93 ± .01 |
| dimer_apo | distal_site_B | 26 | -7.65 | .97 ± 00 |

[a] Schrodinger Glide Gscore

[b] Calculated from PockDrug webserver

**Table S4** Top 10 scoring compounds overall for the dimer interface region

| simulation | region (chain) | rms cluster | Gscore[a] | druggability[b] |
|---|---|---|---|---|
| dimer_apo | interface | 10 | -10.07 | .65 ± .03 |
| dimer_apo | interface | 39 | -9.99 | .83 ± .03 |
| dimer | interface | 15 | -9.72 | .84 ± .05 |
| dimer | interface | 25 | -9.68 | .75 ± .03 |
| dimer_apo | interface | 39 | -9.61 | .83 ± .03 |
| dimer_apo | interface | 39 | -9.58 | .83 ± .03 |
| dimer_apo | interface | 10 | -9.42 | .65 ± .03 |
| dimer_apo | interface | 39 | -9.33 | .83 ± .03 |
| dimer | interface | 6 | -9.3 | .80 ± .02 |
| dimer_apo | interface | 39 | -9.28 | .83 ± .03 |

[a] Schrodinger Glide Gscore

[b] Calculated from PockDrug webserver
