## Supplementary material for "Elucidation of cryptic and allosteric pockets within the SARS-CoV-2 protease": Movie S1

### Slide 1
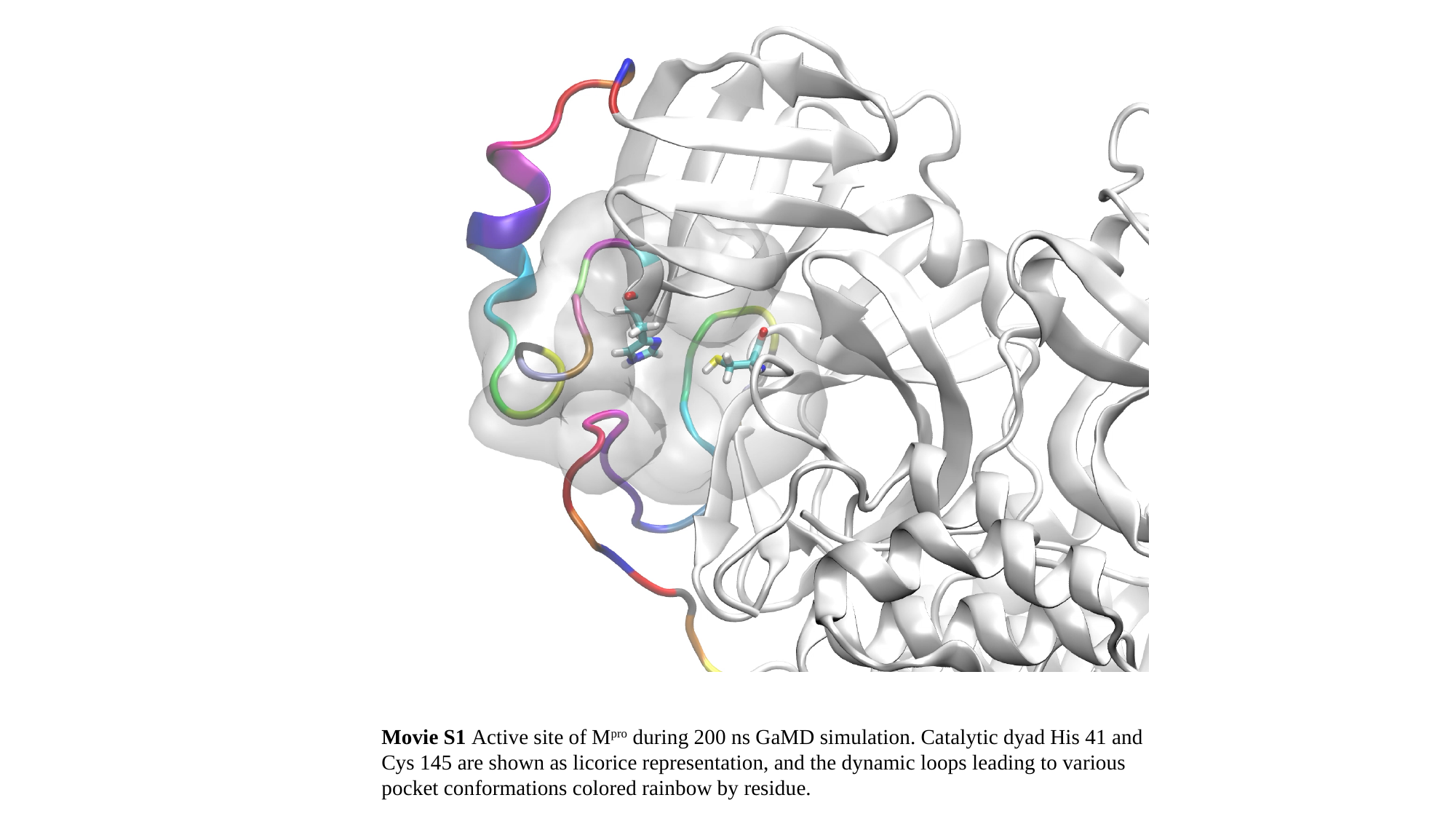

Movie S1 Active site of M^pro^ during 200 ns GaMD simulation. Catalytic dyad His 41 and Cys 145 are shown as licorice representation, and the dynamic loops leading to various pocket conformations colored rainbow by residue.
